# Drug-induced differential culturability in diverse strains of *Mycobacterium tuberculosis*

**DOI:** 10.1101/2024.08.05.606579

**Authors:** Valerie F. A. March, Nino Maghradze, Kakha Mchedlishvili, Teona Avaliani, Rusudan Aspindzelashvili, Zaza Avaliani, Maia Kipiani, Nestani Tukvadze, Levan Jugheli, Selim Bouaouina, Anna Doetsch, Galo A. Goig, Sebastien Gagneux, Sonia Borrell

## Abstract

Differential culturable bacteria grow in liquid culture medium but are unable to form colonies on solid medium. Differentially culturable *Mycobacterium tuberculosis* (Mtb) bacteria, have been found in tuberculosis (TB) patient sputa. We hypothesized that antibiotic treatment can induce differential culturability in Mtb. We investigated the effect of exposure to TB drugs on Mtb culturability using clinical samples from an ongoing TB patient cohort and by conducting several *in vitro* experiments with a diverse set of Mtb strains. In patients, serial sputa were more likely to generate Mtb-positive cultures in liquid as opposed to solid medium, with this liquid culture bias extending up to 5 months post diagnosis. Experimentally, there was a disparity between bacterial time to positivity (TTP) and colony forming units (CFUs) when Mtb was exposed to isoniazid (INH) and rifampicin (RIF) alone or in combination. Cultures recovered from RIF treatment yielded more CFUs on agar plates, but INH-treated cultures had a faster TTP in liquid. Follow up experiments using a fluorescently labelled laboratory strain of Mtb revealed that CFUs overestimated killing by INH treatment. Here we provide evidence in Mtb that drug exposure affects culturability on solid medium, which has implications for treatment monitoring and drug-pathogen interaction studies.

## Introduction

With increasing global concerns over antimicrobial resistance (AMR)^1,2^, antimicrobial stewardship is paramount^3^. Evidence suggests that optimizing therapy in infectious disease patients can help reduce AMR rates^4,5^, which requires adequate treatment monitoring. A key outcome in treatment monitoring is culture conversion; the shift from a positive to a negative bacterial culture from patients undergoing treatment^6,7^. Implicitly, there is a high reliance on culture methods representing all viable bacteria present in patient samples. However, bacterial phenotypes that result in differential culturability can complicate the outcome.

*Mycobacterium tuberculosis* (Mtb) which causes tuberculosis (TB), remains one of the most important bacterial pathogens ^8,9^. TB diagnosis relies on multiple measures taken from patient sputa, including smear microscopy, liquid culture using the BACTEC MGIT 960 system, and solid culture using Middlebrook 7H11 media, among others^10^. These measures, however, are dependent on the bacillary loads in the patient sputa. Furthermore, not all viable bacteria in patient sputa are culturable due to a phenomenon known has differentially culturable tubercle bacteria (DCTB) ^11,12^.

DCTB falls under the umbrella of a general bacterial phenotype known as the viable but non culturable (VBNC) state^13,14^. This has been described in many bacterial species and can be induced *in vitro* through a variety of stressors including hypoxia, starvation and low temperatures^12^.

In Mtb, a predicted 90% of bacteria in patient sputa cannot be cultured^11,15,16^. Current literature has established that for the most part, Mtb preferentially grows in liquid medium, and more of these DCTB can be recovered through supplementation with resuscitation-promoting factors (rpfs) or Mtb culture filtrate (cf)^17^. DCTB can be a complication in treatment monitoring, which, in part relies on culture positivity.

DCTB has been reported in TB patient sputa from the time of diagnosis, up to two months after the advent of treatment^11,18^. In diagnostic cultures, the effect of the host immune response and natural phenotypic variation can be assumed to account for DCTB. The maintenance of DCTB after the advent of treatment, however, allows for the assumption that antibiotic action can contribute to DCTB.

The premise of antibiotic-induced selective viability or differential culturability is plausible based on the target of many antibiotics. Front-line therapy for susceptible TB includes isoniazid, pyrazinamide, rifampicin and ethambutol, which contains mycolic acid, cell wall synthesis, and plasma membrane inhibitors^19^. From this alone, one could posit that prior to sterilization, there could be molecular programs at play that could affect colony formation due to the antibiotic stress response. The question of the impact of antibiotics on culturability or colony formation on Mtb requires attention, as ensuring successful bacterial clearance in patients is key to limit relapse and antimicrobial resistance emergence. Moreover, in drug-pathogen interaction studies, DCTB could allow for the misinterpretation of drug-induced *selective viability* as bacterial *killing*.

In this study, we tested the hypothesis that antibiotics can induce a differentially culturable phenotype in Mtb. We first analysed data from an ongoing TB patient cohort study, and followed this up by experimentally assessing the impact of exposure to the front-line TB drugs isoniazid (INH) and rifampicin (RIF), alone and in combination, on Mtb culturability, using a set of phylogenetically diverse Mtb clinical strains^20^.

## Results

### Liquid culture bias in Mtb clinical samples

We cultured serial clinical samples (n = 449) from 135 Georgian TB patients undergoing treatment, recruited from November 2020 to January 2024. This cohort included 99 multi-drug resistant (MDR-TB) and 36 drug-susceptible cases. For all of these patients, Mtb bacteria were cultured from sputum samples taken at initial diagnosis and during multiple follow-up visits, comprising weekly sampling during the first month, and monthly sampling thereafter. We evaluated preferential culturability in these samples using the BACTEC MGIT 960 system for liquid culture and Middlebrook 7H11 OADC plates for solid culture. From this sampling effort, we surmised the presence of differentially culturable Mtb bacteria, as there was consistently a greater number of positive samples in liquid compared to solid media (p = 0.0023, **Fig 1C**). Moreover, we found that this liquid culture bias persisted up until 5 months post-diagnosis, irrespective of the TB case definition (**Fig 1A & B**).

**Figure 1.**
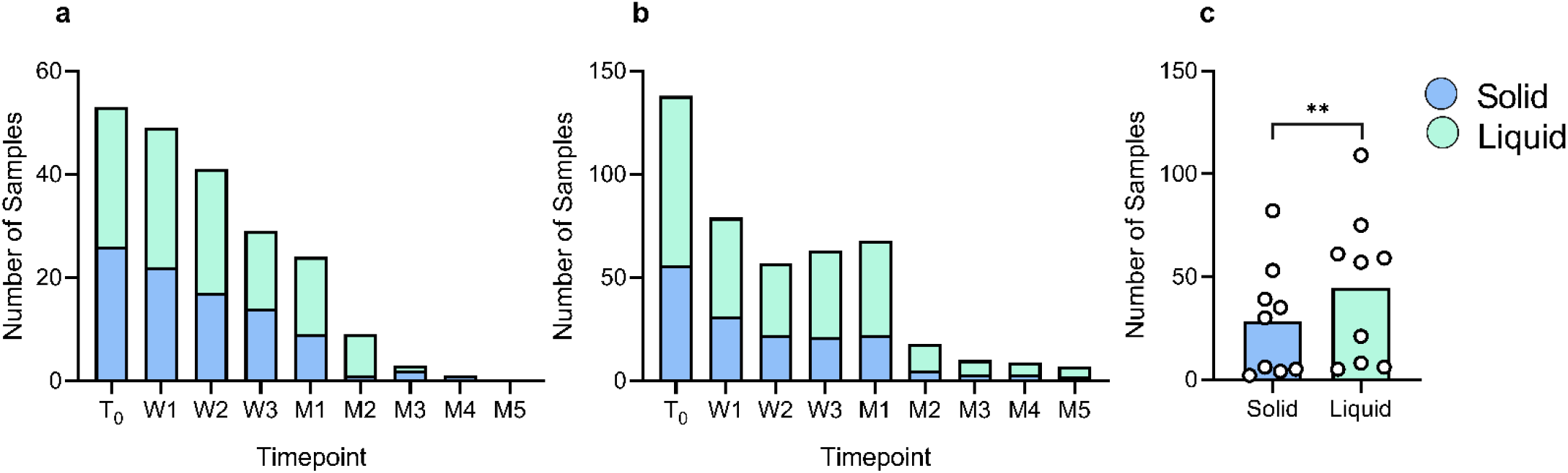
Number of culture positive samples (n = 449) in liquid (green) and solid (blue) media from serial sputa taken from Georgian TB patients (n = 135). Sputa were collected at time of diagnosis (T_0_), weekly (W1-W3), and monthly up to month 5 (M1-M5). (**a**) Number of culture positive samples from MDR-TB patients (n = 99). (**b**) Number of culture positive samples from patients (n = 36) with drug susceptible TB. (**c**) Comparison of total number of all liquid positive (n = 284) and solid positive (n = 165) cultures from November 2020 to January 2024. ** P < 0.01 by paired t test.

### INH treatment results in fewer CFUs, but faster time to positivity in liquid culture

To investigate the contribution of drug exposure to differentially culturable phenotypes in Mtb *in vitro*, we selected a panel of phylogenetically diverse Mtb clinical isolates comprising 3 different lineages (**Table S1**), and subjected them to high-dose short-duration exposure to INH and RIF, alone and in combination. The chosen drug concentrations corresponded to 5X – 180X MIC for all the strains (**Table S1**). After a 48-hour treatment, we removed the antibiotics and quantified the surviving bacterial cells via CFU determination on 7H11 OADC agar plates. We additionally allowed the cultures to recover to mid-log phase by inoculating them into drug-free Middlebrook 7H9 OADC liquid. Our survival assessment indicated significantly less survival in INH-treated cultures compared to RIF alone (**Fig 2A**, p = 0.0033), whereas the combination treatment showed an intermediate level of survival, mainly due to strain-related variation. There was little effect of the Mtb lineage on survival **(Fig S 1&2**). Surprisingly, in monitoring bacterial recovery after treatment, we saw an opposite effect, where INH-treated cultures had the fastest time to positivity (TTP) (**Fig 2B**) compared to RIF alone (p = 0.0123), and in combination with INH (p = 0.0076).To better explore this phenomenon, we turned to metabolic viability assays to disentangle these paradoxical findings.

**Figure 2.**
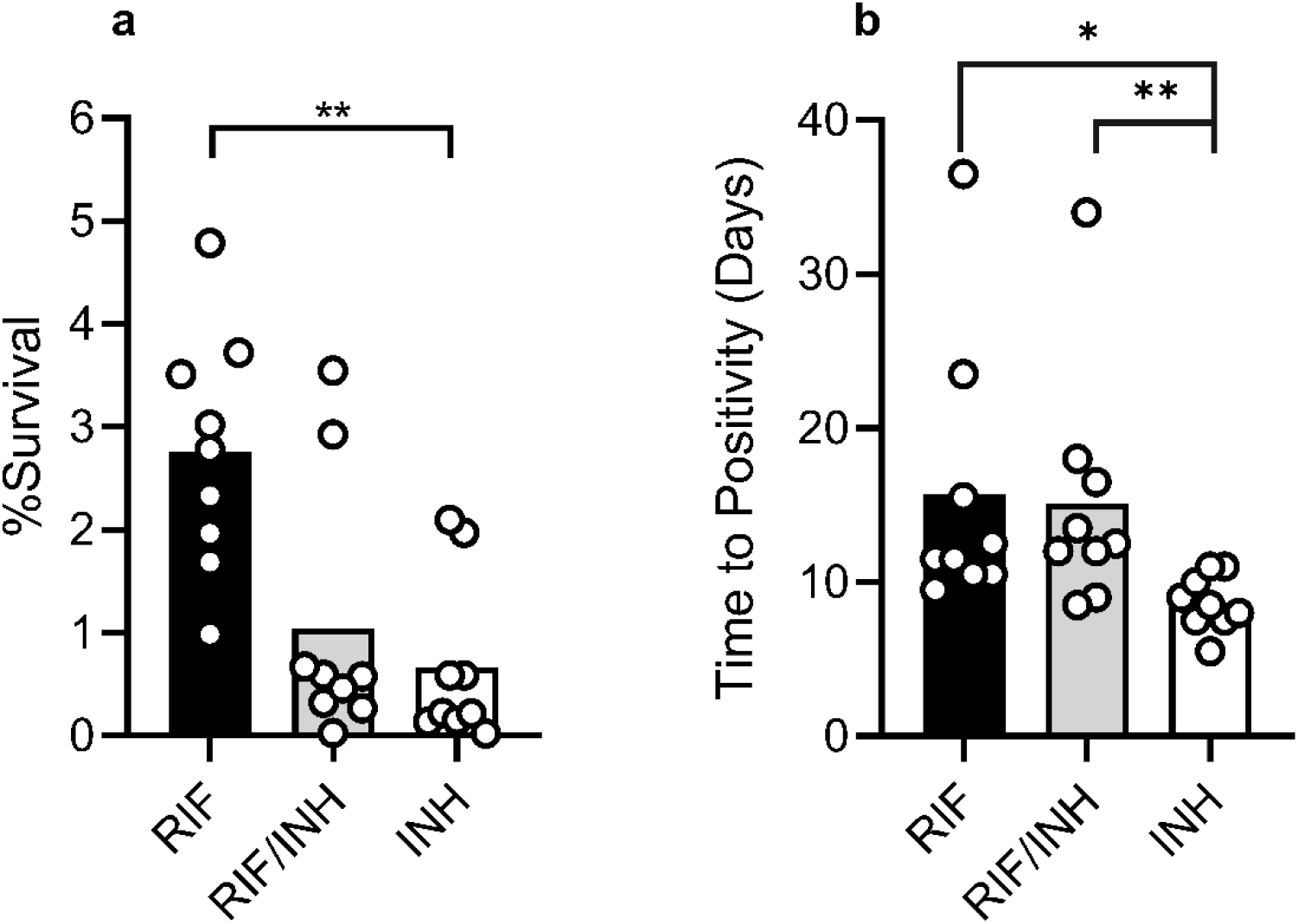
Paradoxical survival and time to positivity in diverse strains of Mtb. Nine Mtb strains (three representative strains each of L1, L2 and L4), were treated for 48 hours with 0.5 μg/ml rifampicin (RIF) and 2 μg/ml isoniazid (INH) alone and in combination (RIF/INH). (**a**) Post treatment survival of strains measured by CFU assay after removal of antibiotics. (**b**) Time to positivity determined by monitoring the duration it took for recovered cultures to reach an OD_600nm_ of at least 0.4 after resuspension in antibiotic-free media. Bars denote mean across strains, each circle represents a mean of n = 2 independent experiment per strain.* P < 0.05, ** P < 0.01 evaluated by Kruskal-Wallis test.

### Metabolic viability assays reveal high metabolic activity in INH-treated bacteria

Looking into the relative ATP output after treatment compared to before treatment, we saw that all treatments led to some increase in the fold-change ATP expression compared to untreated controls (**Fig S4A**). However, the INH-treated cultures showed a much stronger increase in ATP expression (**Fig 3A**). ATP expression after treatment positively correlated with bacterial recovery, which we defined as the inverse of the TTP (**Fig 3C**). ATP expression was also negatively correlated with survival (**Fig S3B)**. In an additional experimental approach, we assessed bacterial viability by the ability of cultures to reduce resazurin to its fluorescent product resorufin. We found that INH-treated cultures had higher levels of resorufin compared to the untreated control (**Fig 3B**) and other conditions (**Fig S4B**). As these results were more a measure of metabolic state than viability, we turned to experiments using fluorescent Mtb.

**Figure 3.**
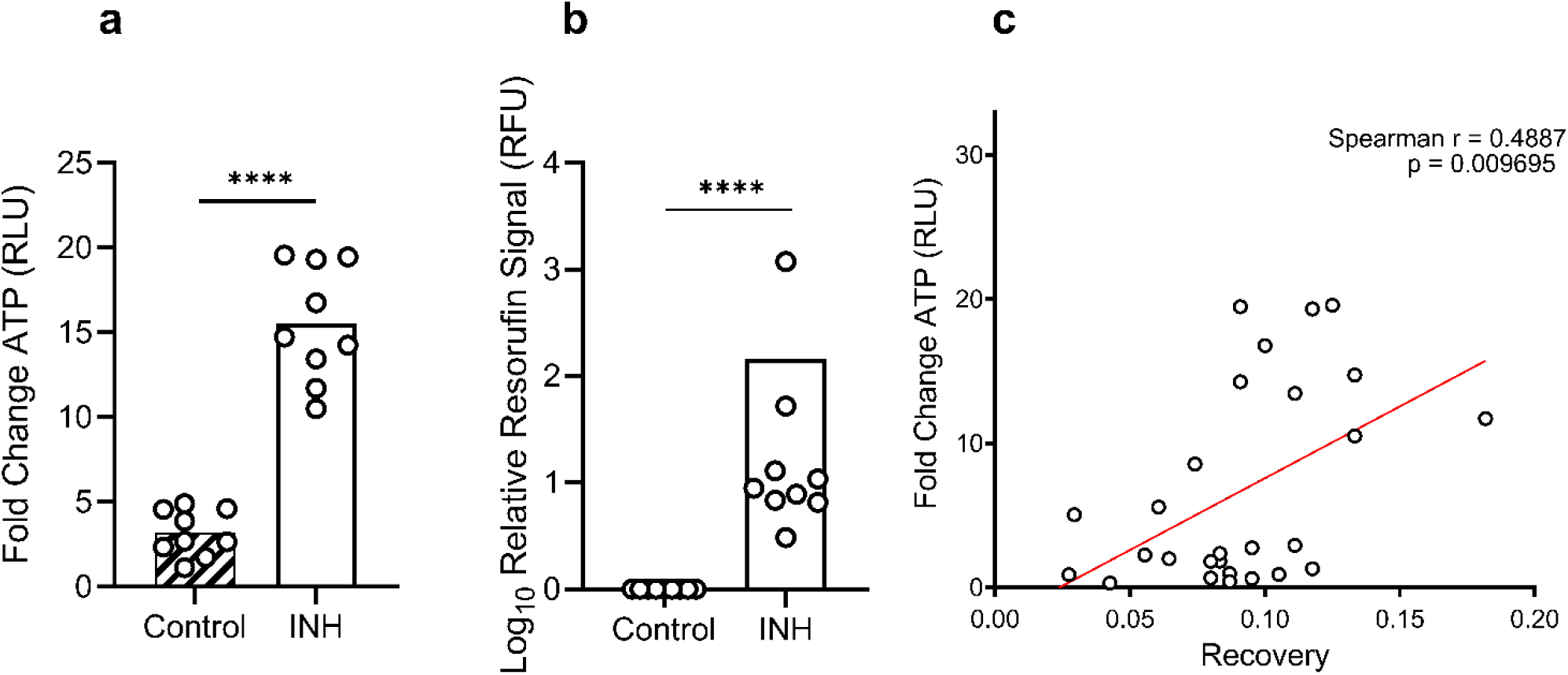
Post-treatment metabolic dysregulation. (**a**) Relative ATP content after INH treatment compared to pre-treatment levels in strains. (**b**) Relative resorufin content as a result of resazurin reduction in cultures treated with INH. (**c**) Spearman correlation of fold change ATP content post-treatment and recovery defined as 1/TTP. **** P < 0.0001 as determined by unpaired t test. Bars denote mean across strains, circles represent mean of n = 2 independent experiments performed in triplicate (a) and one independent experiment in triplicate (b) per strain.

### Fluorescently labelled H37Rv shows no difference in bacterial load across treatments

To further explore the discrepancy between survival and TTP, we repeated the experiment described above, using GFP -labelled H37Rv, the canonical laboratory strain of Mtb. Here, the change in fluorescence after treatment was an additional indicator of the change in bacterial load in each treatment.

In this experiment, there were more CFUs in RIF treated cultures, much like in the diverse strain set, however, cultures recovered from combination treatment or INH treatment yielded far fewer CFUs (**Fig 4A**, p = 0.0185). When looking at TTP, the results mirrored the initial experiment, INH-treated cells had a significantly faster TTP (**Fig 4B**, p = 0.01). INH-treated cultures also had a significantly higher ATP expression compared to the control (**Fig S5**, p = 0.0033). However, we found that the change in GFP fluorescence across all treatments was not significantly different (**Fig 4C**), indicating that INH treatment did not reduce the bacterial population as severely as CFUs would suggest, compared to other treatments.

**Figure 4.**
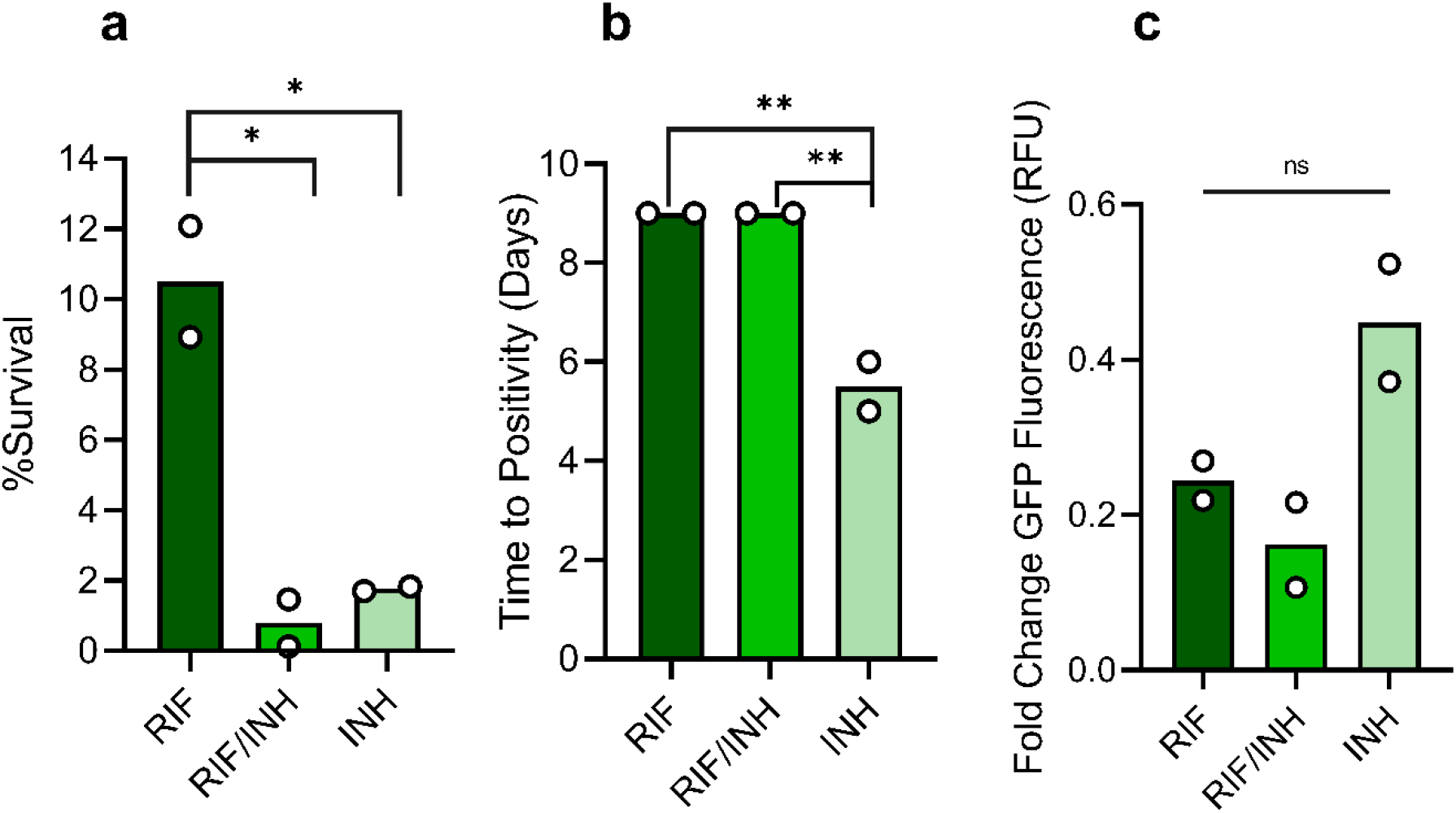
Fluorescently labelled bacteria suggest overestimation of killing by CFU assessment. **(a)** GFP expressing H37Rv survival after treatment with RIF and INH in combination and alone. **(b)** Time to positivity of cultures post treatment. (**c**) Relative GFP fluorescence compared to pre-treatment levels. Each circle represents an independent experiment, bars represent the mean. * P < 0.05 ** P < 0.01 evaluated by one-way ANOVA.

## Discussion

We have shown that clinical Mtb isolated from TB patients undergoing treatment preferentially grow in liquid culture media as opposed to solid, and that this liquid bias extends up to 5 months after initial diagnosis. *In vitro*, we demonstrated that exposure to RIF and INH resulted in a paradoxical culturability, where RIF-treated cultures yielded more CFUs but INH-treated cultures regrew faster in liquid. We have also shown that antibiotic exposure results in high ATP production and that drug-treated bacteria highly reduced resazurin. Finally, using GFP-labelled H37Rv, we demonstrated that the relative decrease in fluorescence across treatments did not significantly differ, despite drug-specific differences in CFUs and time to positivity.

Monitoring culture positivity in TB patients, we found that out of the total of 449 cultures, ∼63.25% were derived from liquid cultures, whereas ∼36.75% were from solid culture, demonstrating a liquid bias. This finding is consistent with previous evidence^18,21-23^. Moreover, this liquid culture bias was maintained up to 5 months post diagnosis. Culture bias is important to consider for cases involving difficult to detect TB such as HIV patients, culture or smear negative TB patients, or in early bactericidal activity investigations where bacterial loads decrease after treatment initiation^24-26^.

Following the hypothesis that exposure to drug per se could lead to this liquid culture bias, we investigated the effect of antibiotic exposure on Mtb culturability on a diverse set of clinical strains that represent globally dominant lineages^20,27^, including those associated with drug resistance^28^. We saw opposite results depending on whether bacteria were exposed to RIF or INH. By CFU determination, there was an average of 3% survival of RIF treatment, and less than 0.5% survival of INH treatment. However, when evaluating recovery in liquid culture, INH-treated cultures recovered almost twice as fast as RIF-treated cultures.

With an outlook to use alternative measures of viability to disentangle our paradoxical CFU and TTP data, we utilized the BacTiter-Glo^29^ and resazurin assays^30^. Where we expected signal loss, all drug conditions had higher signals than the untreated control, with INH treated cells having the highest increase in ATP and largest resorufin fluorescence signal. High ATP after antibiotic exposure has been reported in mycobacteria^31^,which correlated with faster recovery in our work. This contrasts the pattern in untreated cells where high ATP producers grew slightly slower. The resazurin assay is canonically used for drug susceptibility testing in Mtb research, but in our experiments, it seemed to be an indicator of oxidative stress, which has been reported in antibiotic exposed Mtb and VBNCs^13,14,23,32-36^. Combined, we interpreted these data as metabolic perturbation that forms part of the stress response against antibiotics.

In light of these unexpected findings, we repeated our experiments using GFP positive H37Rv. There was no significant difference in GFP fluorescence across all conditions, despite the survival and TTP data corresponding to the initial experiment. As such, we posit that INH-treatment may have induced differential culturability, which affected colony formation. This corresponds with an earlier study done in H37Rv that showed that rpf-dependent DCTB can be triggered with multiple front-line TB drugs^37^. Additionally, antibiotics have been shown to trigger VBNCs in *Staphylococcus aureus*^38^, which further supports our hypothesis.

We note that our work is limited in that, our clinical data is a course-grained assignment of culture bias without quantifying the magnitude of difference in terms of bacterial load, between solid and liquid cultures which others have shown^17,39^. Furthermore, we have not directly shown that DCTB was triggered *in vitro*, however it is a highly plausible explanation for our data.

These limitations notwithstanding, it is clear that CFUs do not always capture the entire viable population despite being a gold-standard measure of viability.

In conclusion, we have presented evidence that antibiotic action can induce a metabolic shift in Mtb that affects colony formation on solid culture, thus supporting a direct contribution of antibiotic stress response in Mtb culturability^14,37,40^. This calls for a careful evaluation of viability assessment when antibiotic exposure is a factor.

## Materials and Methods

### Longitudinal study of clinical Mtb strains

As part of an ongoing prospective cohort study in collaboration with the National Center for Tuberculosis and Lung Disease in Tbilisi, Georgia, we have recruited 135 adult TB patients since 2022. Ethical approval for this study was granted by the Institutional Review Board of the National Centre for Tuberculosis and Lung Disease in Tbilisi, Georgia and the Ethics Commission of North- and Central Switzerland. Individual written consent was obtained from every patient prior to recruitment into the study. The Mtb strains presented in this work were isolated from patient sputum samples up until January 2024. Sputa were collected weekly from the time of diagnosis in the first month, and monthly thereafter, until culture conversion or a change in case definition. After decontamination of sputa, samples were cultured on standard Middlebrook 7H11 agar plates supplemented with OADC and in the BACTEC MGIT 960 culture system in parallel.

### Experimental strain selection and cultivation

Representative strains from a reference set of clinical strains^20^ of lineage 1, 2 and 4 were grown in Middlebrook 7H9 liquid supplemented with 10% ADC (5% bovine albumin-fraction V + 2% dextrose + 0.003% catalase, Sigma-Aldrich), 0.5% glycerol (PanReac AppliedChem) and 0.1% Tween-80 (Sigma-Aldrich), to late lag-phase. These cultures were subsequently plated on BBL^TM^ Middlebrook 7H11 agar (Becton – Dickenson) plates supplemented with oleic acid, albumin, dextrose and catalase (OADC, 0.05% oleic acid (Axonlab) in ADC). Single colonies were picked from each plate and regrown in liquid to establish isogenic cultures, which were subsequently frozen with a final concentration of 5% glycerol. Strains were otherwise routinely cultured in supplemented Middlebrook 7H9 liquid at 37°C shaking at 140 rpm.

### Antibiotic stocks

Antibiotic stocks were prepared from powdered stocks of rifampicin and isoniazid (Sigma-Aldrich), which were diluted in DMSO (AppliChem) such that the final concentration of DMSO in experimental tubes was 0.5%.

### Assessing antibiotic induced differential culturability

#### Experimental set up

Starter cultures of Mtb strains were grown until mid-log phase (OD_600nm_ 0.4 – 0.8) and subsequently calibrated to an OD_600nm_ of 0.05. The calibrated bacterial suspension was divided into four labelled treatment tubes for each strain at a volume of 10 ml. Treatment was subsequently added to each tube (control = no drug, RIF = 0.5 µg/ml, RIF + INH = 0.5 & 2 µg/ml respectively, and INH = 2 µg/ml). Treatment tubes were left shaking at 140 rpm at 37 °C for 48 hours. The initial inoculum was quantified by serial dilution in PBST (Phosphate Buffer Saline + 0.05% Tween80, Sigma-Aldrich) and plated for CFUs on 7H11 OADC plates, which were incubated at 37 °C plates were counted at approximately 21, 28, and 35 days post plating. The initial ATP signal of strains prior to treatment was measured using the BacTiter-Glo assay.

#### Measuring survival and time to positivity

After undergoing treatment for 48 hours, experimental tubes were washed by centrifugation at 1000 x *g*, the resulting supernatant was discarded and pellets were resuspended in fresh Middlebrook 7H9 with OADC. This process was repeated, except the second resuspension was in half the original volume to ensure rapid culture recovery. The washed cultures were sampled for CFU enumeration and ATP determination as described above. Survival was defined as the percentage of CFUs counted after treatment using the baseline pre-treatment CFU count as a basis. Recovering cultures were left shaking at 37 °C at 140 rpm. Growth was monitored to measure time to positivity (TTP), defined as the time at which the culture reached an OD_600nm_ higher than or equal to 0.4.

#### Measuring ATP using the BacTiter-Glo assay

ATP presence was measured in cultures using the BacTiter-Glo assay according to manufacturer’s instructions with an extended incubation time (Promega). Briefly, 75 – 100 μl of BacTiter-Glo reagent was added to white 96 well luminescence plates (Greiner), subsequently an equal volume of culture was added and mixed five times. BacTiter-Glo reagent incubated with an equal volume of Middlebrook 7H9 with OADC was used to control for background luminescence. Luminescence plates were incubated for 45 minutes before luminescence was read using a Tecan Infinite Pro 200 plate reader. Each condition was measured in triplicate. Fold change ATP was expressed as the ratio of the ATP signal post treatment, to the ATP signal in the initial inoculum prior to treatment.

#### Resazurin assay

Washed cultures were inoculated at a volume of 90 µl in triplicate onto clear 96 well plates. 7H9 OADC was also included in triplicate as a negative control. Following this 10 µl of resazurin was immediately added to each well and the plate was incubated overnight.

Fluorescence was measured using the Tecan Infinite Pro 200, with excitation at 560 nm and emission measured at 590 nm. Relative resorufin content was expressed as the ratio of the fluorescence of drug treated signal over the untreated control signal after subtracting background fluorescence.

#### GFP-labelled H37Rv follow up experiments

To complement clinical strain data, the above experiments were repeated using a GFP-positive laboratory strain of Mtb (unpublished), H37Rv, and an RFP-positive H37Rv (unpublished), as a fluorescence background control with the outlook of using the change in fluorescent signal as a measure of bacterial burden. These strains were stably transformed with cassettes constitutively expressing GFP or RFP. Dr. Damien Portevin kindly provided the fluorescently labelled H37rv strains.

#### Fluorescence measurement

Pre and post-treatment GFP signal was measured using the Tecan Infinite Pro 200 plate reader. In brief, 100 μl of culture was added to a black 96 well plate in triplicate. To measure GFP, wells were excited at 435 nm and emission was measured at 535 nm. Background fluorescence was removed by subtracting signal from RFP wells from GFP wells. Post-treatment fluorescence was then expressed as the fold-change in fluorescence compared to pre-treatment measurements.

### Statistical Analysis

Data were visualized and analysed using GraphPad Prism (version 8.2.1), statistical tests used were based on normality of the given data.

## Acknowledgments

This work was supported by the Swiss National Science Foundation (grants 320030-227432, and CRSII5_213514) and the European Research Council (883582-ECOEVODRTB).We also extend our gratitude to Damien Portevin for providing fluorescently labelled H37Rv strains.

## Author Contributions

Study conception: V.F.A.M., L.J., N.T., G.A.G., S.Bor., & S.G ; experimental design: V.F.A.M., L.J., S.Bor., & S.G.; data acquisition and analysis: V.F.A.M., K.M., N.M., A.D., S.Bou. & L.J.; interpretation of data: V.F.A.M., G.A.G., S.Bor., & S.G.; drafted the main manuscript text: V.F.A.M., S.Bor., & S.G.

## Data Availability

All relevant data are within the manuscript and its Supporting Information files.

## Additional Information

Correspondence and requests for materials should be addressed to S.Bor.

