## Supplementary information for "Drug-induced differential culturability in diverse strains of *Mycobacterium tuberculosis*"

**Supplementary Methods**

**Measuring minimum inhibitory concentrations (MIC)**

MICs of parental strains were measured using the REMA method. Briefly, 90 µl of supplemented 7H9 was added to 96 well plates. Then antibiotics were added in a 2-fold dilution series across the plate leaving drug-free wells for controls. Following this 10 µl of bacteria was added in triplicate to a final $OD_{600}$ nm of 0.05. Plates were incubated for 7 days at 37 °C, and then 10 µl of resazurin (Sigma) was added, and left to further incubate overnight. Fluorescence was measured using the Tecan Infinite Pro 200, with excitation at 560 nm and emission measured at 590 nm. Relative growth of each well was calculated by subtracting the background fluorescence of negative control well and expressing growth relative to the signal in drug-free positive control wells. The MIC was defined as the drug concentration at which growth was inhibited by at least 90%.

.

**Table S1:** Strain information

| Parent Strain | Lineage | Sub lineage | MIC INH | MIC RIF | Effective dose INH | Effective dose RIF |
|---|---|---|---|---|---|---|
| **N0069** | 1 | 1.1.1 | 0.025 | 0.1 | 80x | 5x |
| **N0072** | 1 | 1.1.2 | 0.025 | 0.05 | 80x | 10x |
| **N0157** | 1 | 1.2.1 | 0.025 | 0.1 | 80x | 5x |
| **N0052** | 2 | 2.2.2 | 0.0125 | 0.1 | 160x | 5x |
| **N0145** | 2 | 2.2.1.1 | 0.0125 | 0.05 | 160x | 10x |
| **N0155** | 2 | 2.2.1 | 0.0125 | 0.05 | 160x | 10x |
| **N0136** | 4 | 4.3.3 | 0.025 | 0.025 | 80x | 20x |
| **N1216** | 4 | 4.6.2.2 | 0.0125 | 0.025 | 160x | 20x |
| **N1283** | 4 | 4.2.1 | 0.0125 | 0.1 | 160x | 5x |

**Supplementary figures**

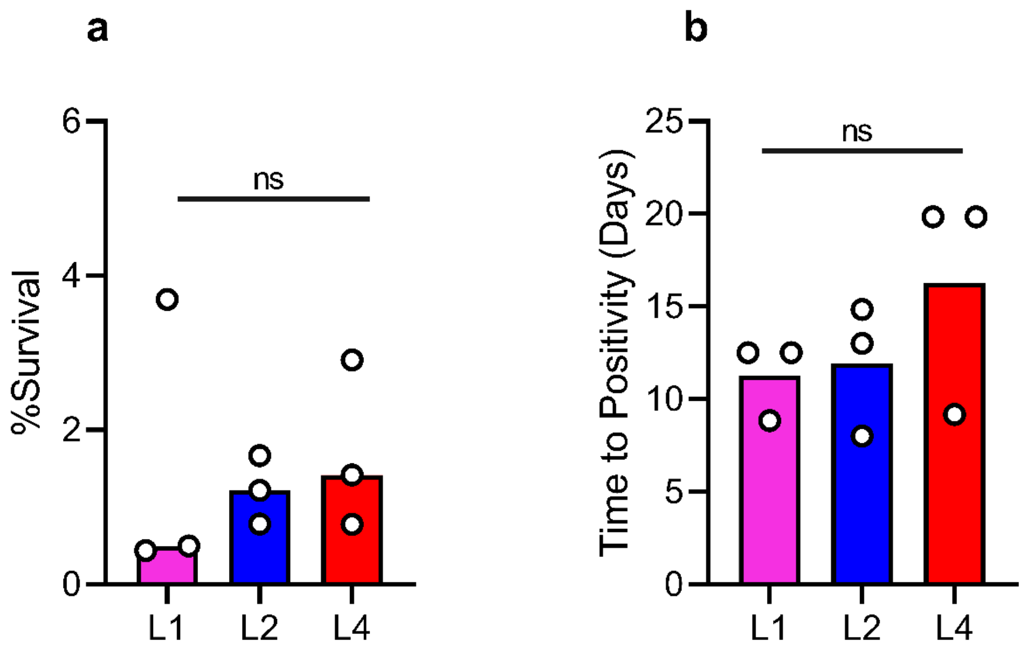


**Figure S1:** Lineage has little effect on treatment and recovery. (**A**) Survival across all treatments by lineage. (**B**) Time to positivity across all treatments by lineage. Bars denote median across lineage, each dot represents mean of n = 2 independent experiments. Statistical differences were assessed by one-way ANOVA.

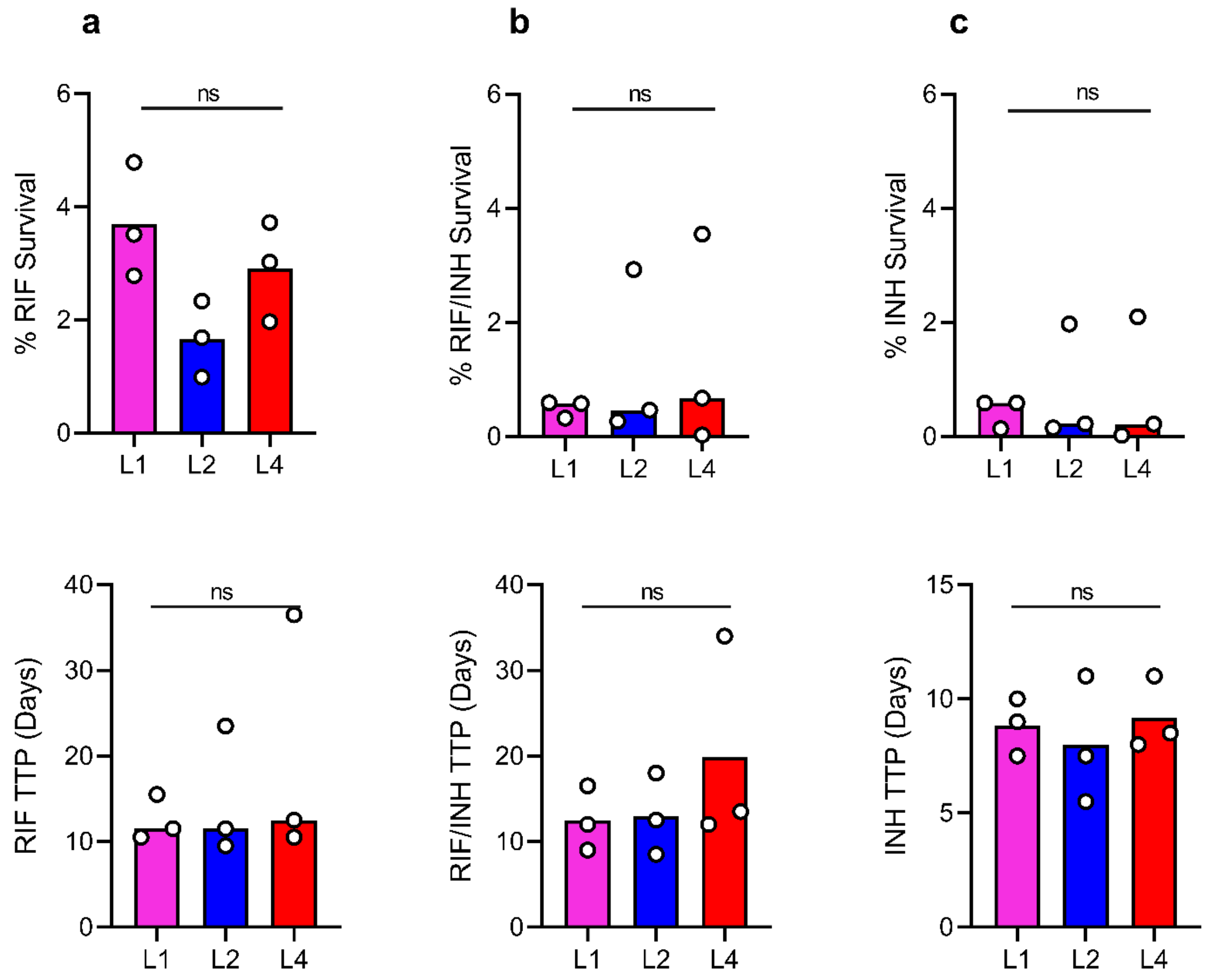


**Figure S2:** Survival (top) and time to positivity (bottom) by lineage, separated by treatment. (**A**) RIF. (**B**) RIF/INH combination (**C**) INH. Bars denote median across lineage, each dot represents mean of n = 2 independent experiments. Statistical differences were assessed by one-way ANOVA.

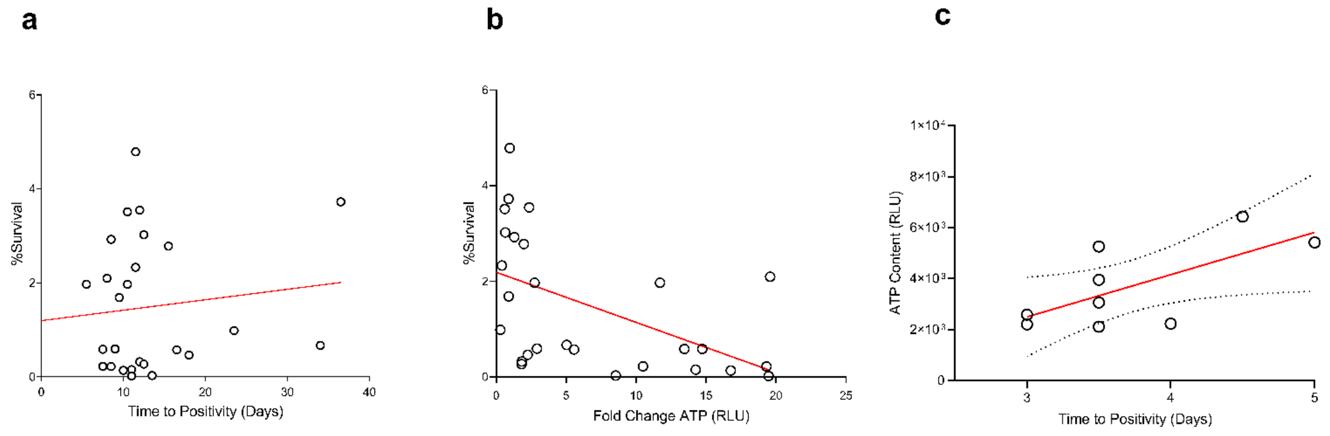


**Figure S3:** Relationship between experimental output parameters. (**A**) Survival vs TTP (Spearman r = 0.1073, p = 0.5943). (**B**) Survival vs Fold change ATP (Spearman r = -0.6160, p = 0.0006243). (**C**) Initial ATP content in untreated cultures vs TTP (Pearson r = 0.6743, p = 00.03146).

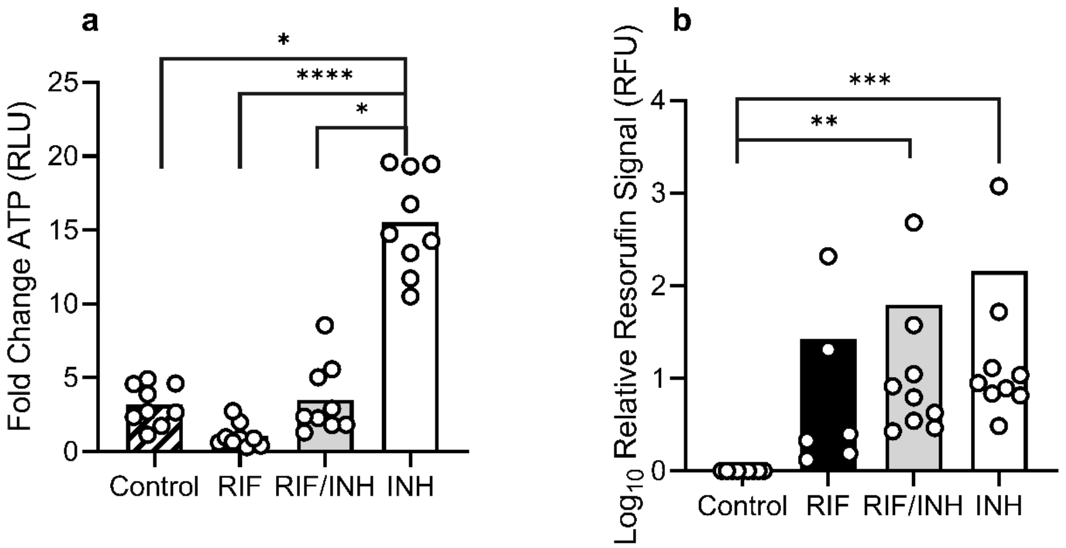


**Figure S4:** Post-treatment energetic and metabolic signatures. (**A**) Fold change ATP across treatments. (**B**) Relative resorufin content across treatments compared to untreated control. Bars denote mean of 9 strains, circles represent mean of n = 2 independent experiments per strain. * $P < 0.05$, ** $P < 0.01$, *** $P < 0.001$, **** $P < 0.00001$ by Kruskal-Wallis.

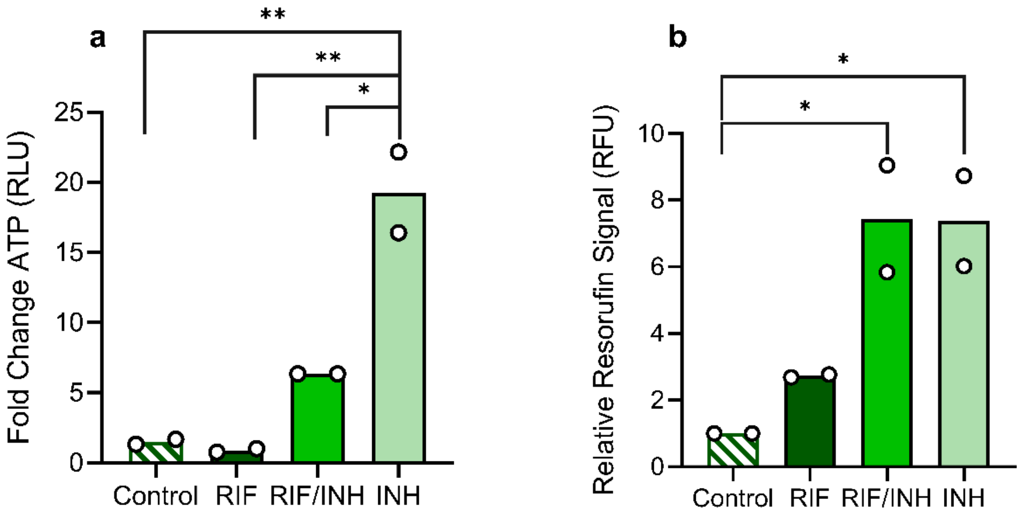


**Figure S5:** Post treatment energetic and metabolic signatures in GFP-labelled H37Rv. (**A**) Fold-change ATP per treatment. (**B**) Relative resorufin content per treatment as compared to untreated control. Data represent mean of n = 2 biological replicates. * P < 0.05, ** P < 0.01 by one-way ANOVA.
